# Neural Networks Are Tuned Near Criticality During a Cognitive Task and Distanced from Criticality In a Psychopharmacological Model of Alzheimer’s Disease

**DOI:** 10.1101/2023.08.16.553626

**Authors:** Forough Habibollahi, Dechuan Sun, Anthony N. Burkitt, Chris French

## Abstract

Dynamical systems exhibit transitions between ordered and disordered states and “criticality” occurs when the system lies at the borderline between these states at which the input is neither strongly damped nor excessively amplified. Impairments in brain function such as dementia or epilepsy could arise from failure of adaptive criticality, and deviation from criticality may be a potential biomarker for cognition-related neurological and psychiatric impairments. Miniscope wide-field calcium imaging of several hundred hippocampal CA1 neurons in freely-behaving mice was studied during rest, a cognitive task of novel object recognition (NOR), and novel object recognition following scopolamine administration that greatly impairs spatial memory encoding. We find that while hippocampal networks exhibit characteristics of a near-critical system at rest, the network activity shifts significantly closer to a critical state when the mice engaged in the NOR task. The dynamics shift away from criticality with impairment of novel object performance due to scopolamine-induced memory impairment. These results support the concept that hippocampal neural networks move closer to criticality when successfully processing increased cognitive load, taking advantage of maximal dynamical range, information content, and transmission that occur in critical regimes.

## 1. Introduction

Recent studies have demonstrated that neuronal populations display complex patterns of electrical activity across a wide spatiotemporal range (He, 2014). Experimental results, spanning from *in vitro* to *in vivo* studies, indicate that these electrical patterns arise spontaneously and are characterized by sequences of activations known as “neuronal avalanches” (Shew and Plenz, 2013). These neuronal avalanches have been studied extensively in the context of criticality, a concept borrowed from physics that describes a state where a system lies in a range between order and disorder. In a near-critical condition, there are long-range connections present between units that produce ordered coordinated events accompanied by some level of disordered fluctuations in the system. This contrasts with a sub-critical system, where there is coordination but no capacity for fluctuations, and also with a super-critical system, where high fluctuations are present but without organized correlations (Chialvo et al., 2010). Over the past decade, critical regimes have been suggested to be a primary mode of neural ensemble activity and also may provide a playing key role in cognitive processes such as learning, memory, and decision-making (Zimmern, 2020). For example, increased cognitive load has been found to help keep the overall brain system in a near-critical state (Zimmern, 2020). Computational studies suggest that near-critical dynamics give rise to increased dynamic range (Deco et al., 2011; Deco and Jirsa, 2012), resulting in an elevated number of cognitive sets and an enhancement of information processing capability (Beggs, 2022). However, it has also been reported that the resting state is associated with near-critical dynamics, whereas cognitive tasks induce subcritical dynamics (Fagerholm et al, 2015; Zimmern, 2020). One possible explanation is that a subcritical system may reduce elements of interference affecting task performance. Thus, whether increased cognitive load shifts neuronal dynamics toward or away from the critical point remains controversial. Understanding this question is crucial, given the potential use of criticality measurement as a biomarker for assessing cognitive function in normal and pathological states such as neurodegenerative conditions (Palutla et al., 2023) and as a potential therapeutic target.

Cortical areas are essential for cognitive functions and processing information, making them the primary targets for studying critical state transitions during cognitive tasks. Other brain regions exhibit varying levels of complexity of task specific neuronal activity, which might influence critical dynamics. The hippocampus is a key brain structure involved in a variety of cognitive processes, particularly memory formation and spatial navigation, but whether these activities involve critical dynamical behaviour has not been previously assessed *in vivo*.

One hallmark of self-organized criticality is that the sizes of neuronal avalanches follow a power law distribution (PLD). This has been observed in numerous *in vitro* studies, including local field potentials (LFPs) of spontaneous activity in acute cortex slices (Beggs and Plenz, 2003; Stewart and Plenz, 2006), and cultured hippocampal neurons (Pu et al., 2013). Additionally, these power law distributions have also been reported in *in vivo* studies, including neocortex of primates (Hoffman and McNaughton, 2002; Petermann et al., 2009), developing cortex of the anesthetized rat (Gireesh and Plenz, 2008), whole-brain of zebrafish (Ponce-Alvarez et al., 2018), and response time in cognitive tasks among healthy adults (Altamura et al., 2012). However, the identification of PLD activity with critical regimes remains uncertain as power-law scaling can also arise in non-critical systems such as random noise activity (Touboul and Destexhe, 2017). Additional metrics such as branching ratio, deviation from criticality coefficient and shape collapse error allow considerably more certainty in identifying critical behaviour (Marshall et al., 2016; Ponce-Alvarez et al., 2018; Ma et al., 2019). Additionally, most of the studies cited above rely on macroscopic variables such as electroencephalograms, LFPs, or serial photographs rather than detecting single neuronal activity and avalanche phenomena. This reliance can increase estimation bias and may lead to incorrect conclusions or skewed analysis results.

Here, we recorded the calcium activity of large populations of single neurons in the mouse hippocampal CA1 area using a miniaturized fluorescence microscope (“miniscope”, Ghosh et al., 2011). We measured criticality characteristics when mice moved freely in their home cage, as well as during the training and testing sessions of a novel object recognition (NOR) task. From widefield imaging of neuronal activity a range of metrics in addition to PLD were used including branching ratio, deviation from criticality coefficient, and shape collapse error. We found that hippocampal network activity was driven closer to the critical point when the animal engaged in the NOR task. Additionally, to assess the impact of a cognitively impaired state on critical dynamics, the observations were repeated after the administration of scopolamine, an amnestic agent that replicates cognitive deficits associated with Alzheimers disease. (San, 2019). We found that scopolamine administration drove the dynamics of the neuronal system away from the critical point. This study enhances our understanding of the role critical dynamics in the task and memory functions of the hippocampal region. We find that increased cognitive load shifts neuronal dynamics toward scale-free behavior near a critical point, while cognitive impairment has the opposite effect.

## 2. Results

Eight C57BL/6 mice were studied while moving freely in their home cage as well as both the training and testing phases of the NOR test (see Supplementary Materials). GCaMP6f AAV (pAAV.Syn.GCaMP6f.WPRE.SV40 virus) was injected into the right hippocampal region followed by gradient index (GRIN) lens implantation. After recovering from the surgeries, a metal baseplate used to support the miniscope device was attached to the animals’ heads to which a UCLA miniscope V3 was attached during experiments connected to a computer interface board allowing data acquisition via a coaxial cable during recording sessions. We recorded from populations of 290 ± 42 CA1 neurons in 15-minute recording sessions. Calcium events associated with the detected neurons were derived from the recordings and used to identify neuronal avalanches from the population activity (Figure 1). Neuronal avalanches were defined to start and stop when the population activity (i.e., the total number of contributing events) surpassed a certain threshold of the entire network activity. These avalanches were then employed to identify the criteria for criticality. Firstly, scale-free dynamics in both temporal and spatial domains and power-law distributions of observing avalanches with a certain size (S) (number of contributing events) and duration (D) (time) are the hallmark of criticality. Second, in critical systems, the power-law exponents fit to avalanche size (τ) and duration (α) are related by a third predicted exponent as 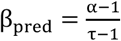. Employing the exponent relation between S and D near-criticality, 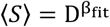, the mean avalanche size (〈*S*〉) observed at a given duration can be predicted using α and τ. When 〈*S*〉 is plotted against avalanche duration, the difference between the empirically derived β_fit_ and the predicted β_pred_ is termed the Deviation from Criticality Coefficient (DCC) which provides another strong metric of criticality, being minimized near the critical state. Branching Ratio (BR) is the third measure and demonstrates the ratio between the number of active units at time t + 1 to the number of active units at time t which is tuned near 1 at criticality. The error from the Shape Collapse analysis (SC error) is the final metric of criticality to demonstrate how accurately avalanches of all profiles can be collapsed to the same universal shape. The error between data and the universal scaling function is minimized at criticality (see Supplementary Materials, Marshall et al., 2016; Wilting and Priesemann, 2018; Ma et al., 2019; Spitzner et al., 2021 for details). With these measures, we sought to evaluate whether a memory-related cognitive task would move the network dynamics closer to criticality compared to resting state activity recorded in the home cage.

**Figure 1.**
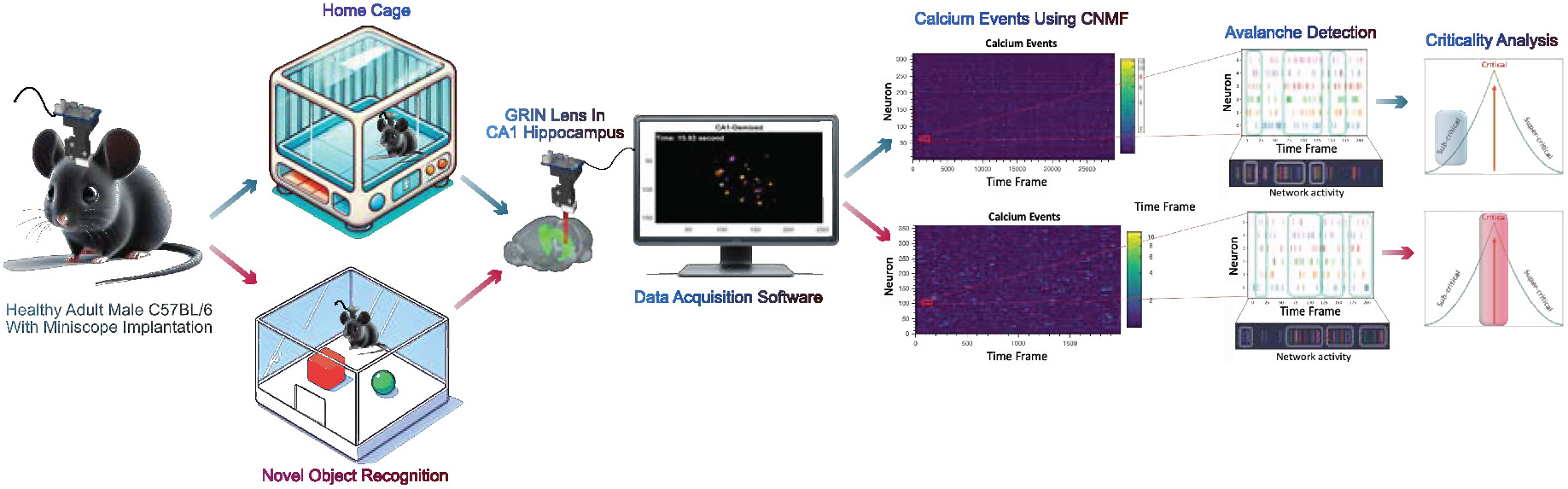
Schematic overview of the study. A GRIN lens was implanted into the hippocampal CA1 area to observe the calcium activity of GCaMP6 virus-labelled neurons in healthy adult male C57BL/6 mice. The animals were then recorded both in their home cage and during the Novel Object Recognition (NOR) task. Minian analysis pipeline was used to identify spatial footprints as well as the temporal traces of detected neurons in each recording session. The raw calcium activity was deconvolved to extract the calcium event signal which were then used to extract neuronal avalanches for the criticality analysis. The criticality-based metrics were then compared between home cage and NOR recordings.

### 2.1 Hippocampus neuronal networks are tuned near-criticality during a cognitive task

We compared critical metrics recorded during free cage exploration and when the animals were in the testing phase of the NOR (see Supplementary Materials). The NOR experiment evaluates the recognition memory of the animals and particularly involves their spatial memory performance. Hence, it applies an increased cognitive load while the animals are exploring both novel and familiar objects actively utilizing spatial memory. We found that when mice in the process of successfully distinguishing between the novel and the familiar object, neural activity shifted closer to a critical state compared with the resting state recording in the home cage.

We used 8 healthy adult male C57BL/6 mice in recording sessions of 15 minutes in their home cage as well as both phases of the NOR test (Figure 2.a). These animals are then exposed to the experimental arena for habituation (see Supplementary Materials) before the familiarization phase (two similar objects) and the testing phase (one familiar and one novel object) of the NOR experiment (Figure 2.b). The animal trails during these phases as well as the object locations in the arena are recorded and drawn by the top-view-based behavior analysis system used in this work (TopScan: TopView Analyzing System 2.0; Clever Sys Inc). A sample trail which is shown in Figure 2.c outlines the higher amount of exploration time around the novel (red) object in the testing phase. The behavioural results showed that on average, the mice spent a longer period of time exploring the novel object in the testing phase while no significant difference was detected between the exploration times of the two similar objects in the familiarization phase (see Figure 2.d).

**Figure 2.**
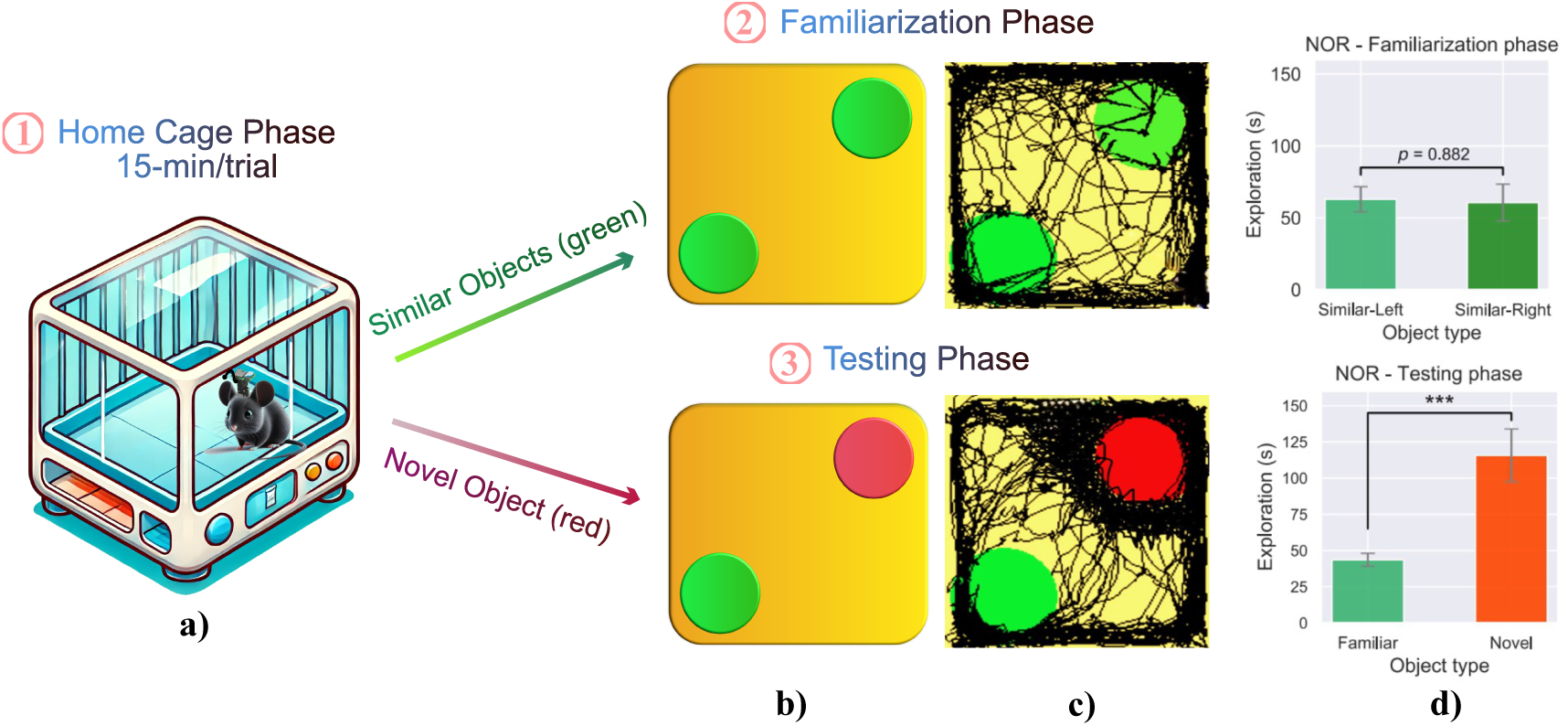
NOR experimental setup and behavioural performance of the mice. **a)** A mouse with the miniscope device mounted on its head is recorded in its home cage. **b)** The setting of both phases in the NOR experiment; top: familiarization phase with 2 similar objects inside the arena, bottom: the testing phase where one of the familiar objects is replaced with a novel one. **c)** Sample mouse trails in black recorded using the TopScan software in the top: familiarization and bottom: testing phase of the NOR experiment. The novel object in the testing phase is colored red. **d)** The average exploration time around each of the objects in the top: familiarization and bottom: testing phase. In the familiarization phase, there was no significant difference detected between the average time spent exploring each of the objects. However, the exploration time of the novel object in the testing phase was significantly higher than the familiar object (One-way ANOVA test, *** p−value < 0.005). Error bars, SEM.

The miniscope recordings were processed using the Constrained Non-negative Matrix Factorization (CNMF) algorithm (Zhou et al., 2018) and the spatial footprints of the detected neurons, as well as the corresponding calcium events, were extracted (see Supplementary Materials for more details). Using these calcium events, the activity of each neuron was binarized in each recorded frame (i.e., 33.3 ms) was binarized, based on being active or not, i.e., 1 and 0. Neuronal avalanches were then calculated as cascades of events with initiation and termination determined after reaching a certain threshold of network activity (i.e. the sum of all recorded activity within the frame). The analysis was robust to changes in threshold over a range of 30 to 70 % of network activity. These avalanches were then utilized in extracting the criticality metrics. Figures 3.a-b compare the power-law distributions and the DCC measures obtained from one of the animals in its home cage and the testing phase of the NOR experiment. While power-law distributions could be fitted to the avalanche size and duration in both groups of recordings (α = 1.937 ± 0.070, τ = 2.597 ± 0.091 in NOR experiments (testing phase); and α = 1.540 ± 0.056, τ = 2.367 ± 0.100 during cage recordings), the lower DCC measure in Figure 3.b indicates the system operating closer to criticality. Figures 3.c-e illustrate the distribution of the criticality metrics in different recording sessions in the cage or the testing phase of NOR (NOR: indicates the testing phase of the experiment in the figures of this article). Comparison of the NOR (colored in cyan) and Cage (colored in orange) sessions shows the shift of the CA1 network dynamics toward criticality during the task-present sessions and the testing phase of NOR.

**Figure 3.**
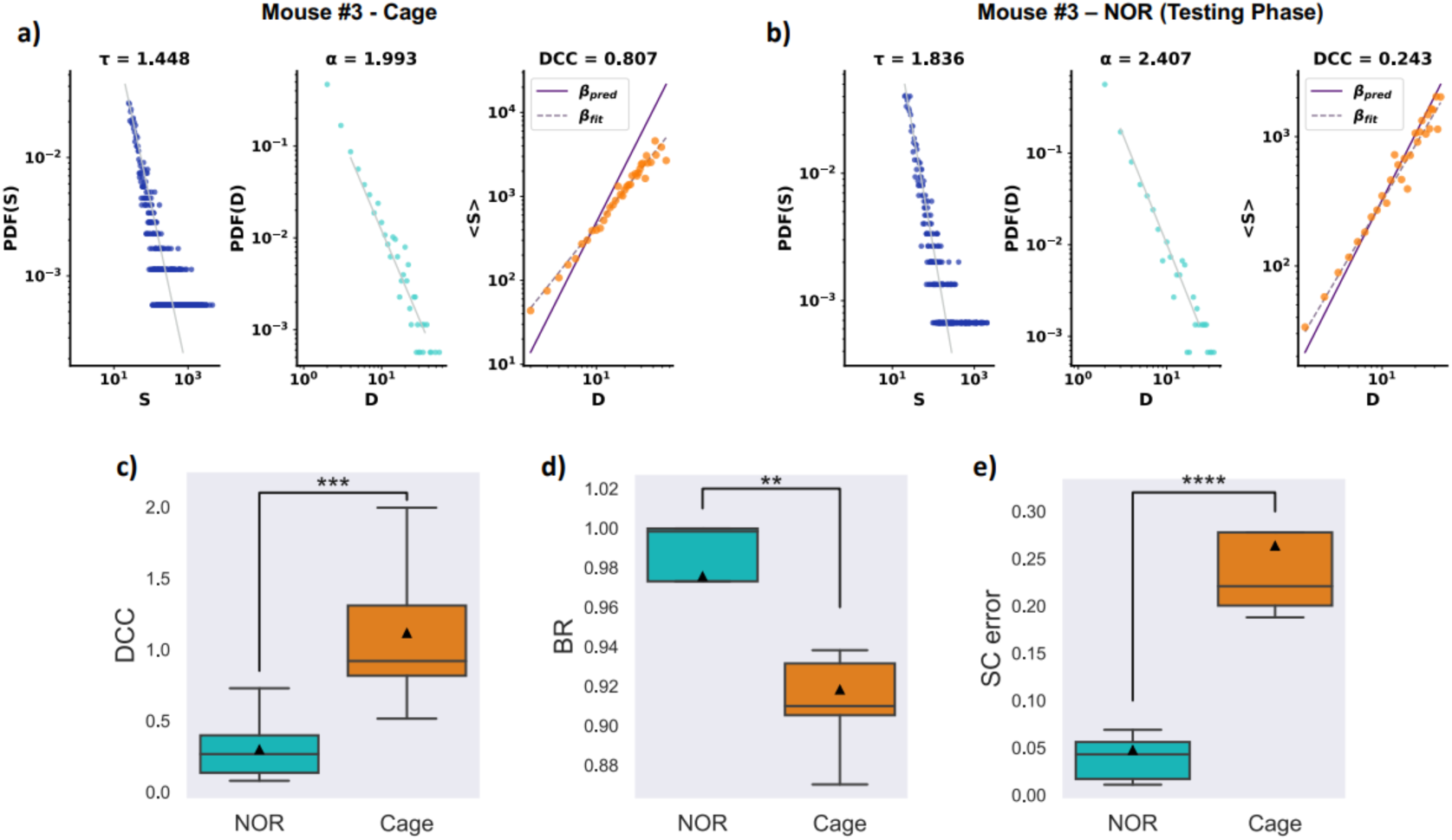
Criticality arises as the animals are engaged in a cognitively demanding task, NOR. Avalanches are quantified as a function of the number of contributing calcium events (S) from all neurons in the region of interest and the duration of the avalanche (D). The Probability Distribution Function (PDF) for detecting avalanches of a specific size (blue) or duration (cyan) can be fit with power-laws, resulting in the generation of two exponents (τ, α) respectively; in the log-log plot, the exponent represents the slope of the line. In critical systems, the following relation holds between the size and duration exponents: 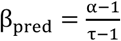. Deviation from criticality as measured by the Deviation from Criticality Coefficient (DCC), is the quantitative difference between the predicted β_pred_ exponent (solid purple line) and the observed β_fit_ exponent (dashed grey line). **a)** The PDF fitted to avalanche size and duration and the DCC measure in the recordings from an example mouse inside the home cage. **b)** The PDF fitted to avalanche size and duration and the DCC measure in the recordings from an example mouse during the testing phase of the NOR experiment. **c, d, e)** DCC, BR, and SC errors were extracted for all the recordings and compared between Cage and the testing phase of the NOR experiment. The illustrated trend in all measures supports the conclusion that the system tunes near-criticality during a cognition task such as NOR. The NOR recordings display DCC and SC error values significantly closer to 0 and branching ratios significantly closer to 1; features that are missing in the cage recordings. (One-way ANOVA test, **** p−value < 0.0005, *** p−value < 0.005, ** p−value < 0.05).

### 2.2 Scopolamine-induced memory impairment drives system dynamics away from criticality

The neurotransmitter acetylcholine is involved in the acquisition and encoding of new information (Hasselmo, 2006). Additionally, it has been observed that the administration of acetylcholine muscarinic receptor antagonists such as scopolamine can significantly impair recognition memory (Tinsley et al., 2011; Kowalczyk et al., 2020). To further evaluate the potential roles of criticality-tuned dynamics in recognition memory, one month after the completion of NOR experiments, the animals were employed in another set of NOR experiments. At this stage, 2 mice were removed from the study due to showing signs of reduced mobility and difficulty mounting miniscope devices. In the first week, the animals were recorded after injection of saline (0.9% NaCl, intraperitoneal, i.p.; 10 mL/kg body mass), 20 minutes before the familiarization phase of NOR (Tinsley et al., 2011). After a 2-week rest interval, the same 6 animals were injected with scopolamine (intraperitoneal, i.p.; 0.6 mg/kg body mass) 20 minutes before the first phase of NOR. The performance of the animals was then compared among the two stages of the study to evaluate the effects of scopolamine-induced memory deficits on the criticality of the hippocampus CA1 network. The number of detected neurons was 284 ± 14 which was not significantly different from the pre-injection levels (p = 0.936, One-way ANOVA test). The average firing rate of the population showed a non-significant reduction in the trials post scopolamine administration compared to the saline-injected control group (3.218 ± 0.839 Hz and 2.932 ± 0.712 Hz, saline and scopolamine administration respectively; p = 0.563, One-way ANOVA test).

First, to confirm the scopolamine impairment of NOR performance, we measured the discrimination index, the global habituation index, the recognition index, and the preference index (Antunes and Biala, 2012) to evaluate the object recognition memory during the NOR test. The discrimination Index (DI) allows discrimination between the two objects in the testing phase of the experiment. DI uses the difference in exploration time for the novel and familiar objects, dividing it by the total amount of exploration of both objects. The score obtained from this test ranges from −1 to +1, where a positive score indicates that the subject spent more time with the novel object, a negative score indicates that the subject spent more time with the familiar object, and a score of zero indicates no preference. The global habituation index (GI) is calculated by dividing the total time spent exploring the two objects during the familiarization phase by the total time spent exploring them during the testing phase. GI tends to be lower than one in healthy animals. The Recognition Index (RI) is calculated as the time spent exploring the novel object divided by the total time spent exploring both objects during the testing phase and is considered the primary measure of retention (Schindler et al., 2010). Therefore, an RI index above 0.5 indicates novel object preference, and below 0.5 shows familiar object preference. Finally, the preference index (PI) is calculated as the ratio of the time spent exploring one of the two objects during the familiarization phase to the total time spent exploring both objects. Therefore, a preference index above or below 0.5 indicates object preference and a value close to 0.5 shows no preference.

The results in Figure 4 compare the NOR experiments following saline (Figure 4.a-b) and scopolamine (Figure 4.c-d) injections. While there is a significant difference between the exploration times of the familiar and novel objects in the testing phase of saline experiments (Figure 4.b), this difference disappears after scopolamine injection (Figure 4.d). Additional evidence of scopolamine impairing the recognition memory of the mice was found using the introduced indices of DI, GI, RI, and PI, described above shown in Figure 4.e. Compared to the saline control group, scopolamine injection significantly reduced the ability to discriminate the novel object (DI), increased the ratio of the global exploration time in the familiarization phase to the testing phase (GI), decreased the time spent exploring the novel object relative to the total object investigation time in the testing phase (RI), and kept the preference index between any of the two similar objects in the familiarization phase near 0.5, similar to saline controls. These preliminary experiments verified previous reports of scopolamine-induced deficits in NOR performance in our cohort of animals with miniscopes.

**Figure 4.**
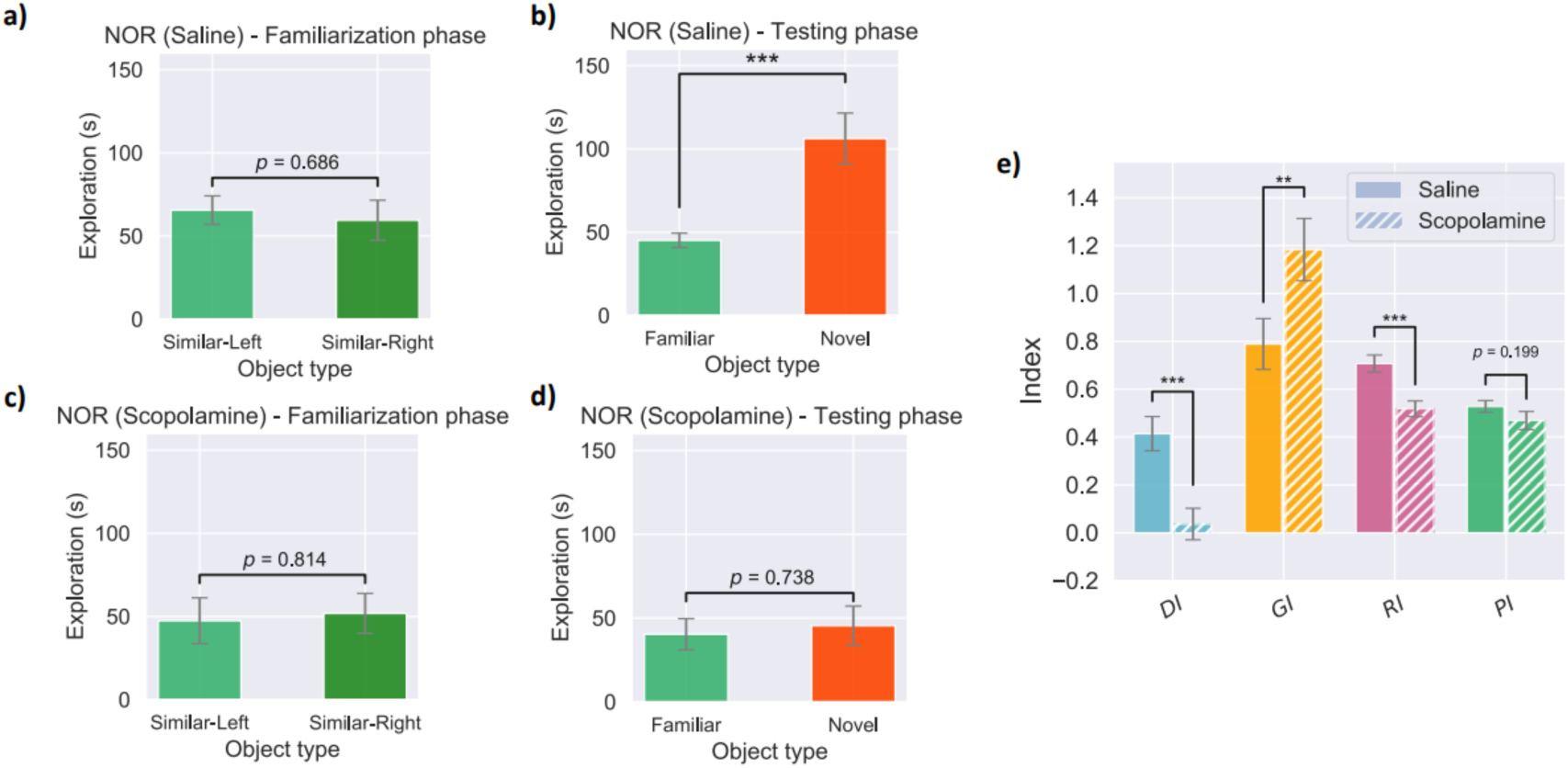
Scopolamine administration impairs spatial memory and disrupts the mice’s performance in the NOR testing phase compared to saline-injected control experiments. **a, b)** Saline control NOR experiments. The average exploration time around each of the objects in the a) familiarization and b) testing phase. In the familiarization phase, there was no significant difference detected between the average time spent exploring each of the objects while in the testing phase, a significantly higher average exploration time was detected around the novel object (One-way ANOVA, *** p−value < 0.005). **c, d)** Scopolamine injected NOR experiments. The average exploration time around each of the objects in the **c)** familiarization and **d)** testing phase. Scopolamine impairs spatial memory affecting the animals’ performance in the NOR test. There was no significant difference observed between the average exploration time of the animals around the familiar and the novel object in the testing phase after scopolamine injection (One-way ANOVA test, p−value = 0.738). e) Comparing the DI, GI, RI, and PI indices in two groups of saline and scopolamine-injected animals. The significant decrease in the discrimination and recognition indices (One-way ANOVA, *** p−value < 0.005), as well as the significant increase in the global habituation index (One-way ANOVA, ** p−value < 0.05) after the scopolamine administration all, indicate spatial and recognition memory deficits induced by scopolamine. Error bars, SEM.

Intriguingly, the functional deficits induced by scopolamine changed the dynamics of the system which was driven further from criticality compared to saline controls (see Figure 5). We repeated the comparison of the three criticality metrics (i.e. DCC, BR, and SC error) for the NOR (Saline), NOR (Scopolamine), and Cage groups. The results confirm our hypothesis that the NOR (Saline) group has a significantly lower average DCC and SC Error while showing a significantly higher BR measure on average. The statistical significance of these findings was evaluated using a pairwise post hoc Tukey’s test and the results are illustrated in Figure 5.

**Figure 5.**
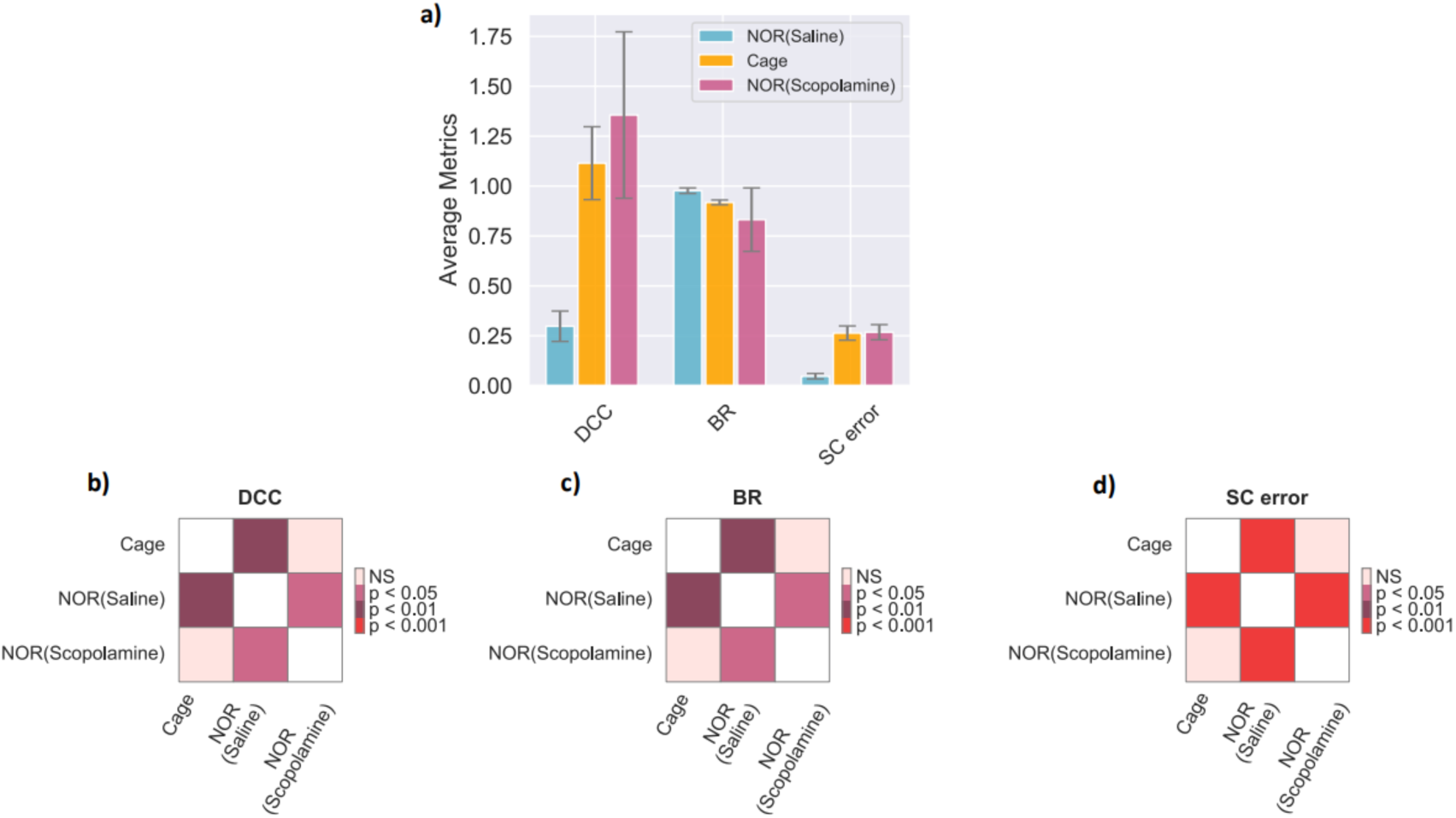
The recognition and spatial memory deficits induced by scopolamine injection tune the network away from criticality compared to saline controls showing dynamics more similar to rest periods in the home cage. **a)** Comparison of the average extracted metrics of criticality in the cage, the testing phase of NOR(Saline), and the testing phase of NOR(Scopolamine) experiments. In all three measurements, the NOR(Scopolamine) test indicates a system that is significantly closer to a critical state compared to both the cage and NOR(Saline) groups. Error bars, SEM. **b, c, d)** The significance matrix using pairwise post hoc Tukey’s test. A significant difference is detected in all three metrics between cage and NOR(Saline) as well as NOR(Scopolamine) and NOR(Saline). There is no significant difference detected between cage and NOR(Scopolamine) groups in any of the measures indicating that both systems are tuned far from criticality.

## 3. Discussion

In this study, we recorded the calcium activity of hippocampal CA1 neuronal populations in two groups of mice: healthy mice and mice with cognitive impairment acutely induced by scopolamine. We analysed and compared their hippocampal criticality characteristics while they moved in their home cage and performed a novel object recognition (NOR) task. The findings demonstrate that cognitive task engagement is associated with enhanced critical dynamics, manifested as a decrease in the deviation from criticality coefficient (DCC) and shape collapse error (SC error), as well as an increase in the branching ratio (BR). Conversely, pharmacological impairment of cognitive function induced by scopolamine produces the opposite effect.

A range of theoretical and experimental findings have suggested a link between critical dynamics in the brain and cognitive activity (Sreekumar et al., 2014; Nielson et al., 2015). From theoretical considerations, if the brain operates in a critical state, critical dynamics will facilitate transmission of information, enhance neural information content, and provide a wide dynamical range of cognitive activity. Additionally this regime will maximize the amount of information encoded and processed by neural circuits, enabling the network to adapt to a wide range of inputs and environmental changes. Our results that increased cognitive load drives neuronal dynamics toward the critical point are very consistent with this hypothesis. On the other hand, previous studies have also found near-critical dynamics in the resting state and sub-critical dynamics during tasks (Beggs and Plenz, 2003; Fagerholm et al, 2015; Cocchi et al., 2017; Zimmern, 2020), suggesting that a subcritical system may decrease interfering factors affecting task performance. We suspect this discordance results from subsampling of the data, as these studies depend on macroscopic variables like EEG or LFP rather than individual neuronal activity to quantify criticality characteristics, which increases estimation bias and may lead to incorrect conclusions or skewed analysis results. Another possible reason may be regional brain area activity, as different brain areas might display varying levels of complexity in neuronal activity for individual tasks. These variances may significantly impact the dynamics of criticality.

An interesting and potentially contentious question related to the current study is whether brain dynamics are situated exactly at criticality or near-critical states. Some studies find dynamics situated precisely at the critical point (Beggs and Plenz, 2003; Bellay et al., 2015; Jannesari et al., 2020), while others, mostly from in vivo spiking studies, report brain dynamics residing in a sub-critical region (Hahn et al., 2010; Ribeiro et al., 2010; Priesemann et al., 2013, 2014; Ma et al., 2019; Wilting and Priesemann, 2019) similar to our findings. The presence of a sub-critical brain state has some interesting implications. A near-critical state possesses computational properties similar to criticality but with a slight reduction in other computational properties associated with the adjacent sub-critical phase. This is because the critical region is ‘stretched’ in a finite and input-driven system like the brain (Wilting and Priesemann, 2018). It therefore still enhances information processing, but being at a certain distance from the critical point could reduce the incidence of dysfunctional neural behaviour, of which abnormal oscillatory activity such as seizures are a possible manifestation (Priesemann et al., 2014). Nevertheless, another conceptual framework has been recently proposed, classifying brain dynamics as “quasicritical” (Beggs, 2022). This proposes that due to the consistently present external drive applied to all functional neuronal ensembles, the fully inactive phase for BR< 1 is no longer present, and thereby, we no longer observe a phase transition between active and inactive states. Due to the absence of this phase transition, there is also no longer a “critical” point. The presence of an external drive prevents the cortex from reaching a critical phase transition point and based on available data, it is expected that such an external drive will always be present. Instead, this type of dynamics is referred to as “quasicritical”. Whether the findings presented here are interpreted as slightly sub-critical or quasicritical, we found that there was a transition towards a critical regime with increased cognitive load. We suspect that this will be generally observed with increased cognitive loads, and it will be of interest to see if this is confirmed in a range of cognitive tasks. Earlier studies identified critical dynamics with power law observables of both event lifetime and spatial distributions. Evidence of these power-law observables has been found in a wide range of *in vitro* (Beggs and Plenz, 2003; Friedman et al., 2012; Habibollahi et al., 2023) and *in vivo* studies (Gautam et al., 2015; Jannesari et al., 2020), utilizing different recording methods (Priesemann et al., 2014; Fagerholm et al., 2015). Nonetheless, it has become apparent that these measures are necessary but not sufficient to identify critical dynamics, since many noncritical processes also exhibit power laws (Priesemann and Shriki, 2018). Consequently, several more rigorous metrics have been developed and recently implemented like the deviation from criticality coefficient, the shape collapse error, and the branching ratio (Sethna et al., 2001; Friedman et al., 2012; Marshall et al., 2016; Wilting and Priesemann, 2018; Ma et al., 2019). All these metrics have been utilized in the current study to better characterize the critical dynamics of the system, thereby providing more reliable results than relying solely on power-law observables.

The miniscope technique, in principle, provides significant advantages for observing critical dynamics behaviours in freely moving animals. Primarily, it allows a large number of neurons comprising a substantial proportion of the activated network to be recorded over considerable periods of time (Ghosh et al., 2011). However, calcium imaging does not achieve the same temporal resolution as electrophysiological recording, due to the slow dynamics of calcium activity and fluorophores. Nonetheless, we find this method is suitable for this study as it allows single action potentials to be detected at low frequencies, within the firing rates typically observed in cognitive tasks with electrode recordings (Power and Sah, 2002; Harding et al., 2020). Miniscopes in mouse hippocampus have been used extensively to record place cell behaviour (Sun et al., 2021; Wirtshafter and Disterhoft, 2022), phenomena generated by complex information encoding and decoding (Sun et al., 2024), with similar results to electrophysiological recording, suggesting a high level of functional correlation between miniscope and electrophysiological methods. Additionally, calcium signals from multiphoton imaging have been used as the basis for a brain-computer interface, again demonstrating a close relationship between calcium transients and “ground truth” electrophysiological signals (Clancy et al., 2014; Trautmann et al., 2021; Sun et al., 2023).

The premise that optimal brain function correlates with critical dynamical behavior has prompted numerous studies to associate a spectrum of cognitive pathological conditions, including epilepsy, schizophrenia, major depressive disorders, and Parkinson’s disease, with suboptimal critical behavior (Zimmern, 2020). Despite some identified associations, these investigations are frequently constrained by their reliance on inferring critical behavior solely through power law calculations of potentially relevant variables. Our present study contributes to validating the observation of these dynamics by employing more robust metrics at the level of neuronal ensembles in a pharmacological mouse model of neurodegenerative diseases induced by scopolamine. Scopolamine treatment was used in the study because it has well-described effects on memory and has been considered as a model for neurodegenerative conditions such as Alzheimer’s disease, in which cholinergic neurotransmission to the hippocampus is known to be impaired (San Tang, 2019; Kowalczyk et al., 2020). We hypothesized that the cognitive degradation would be accompanied by changes in criticality measures, reflecting the impaired functional state of the system. The results presented in this study support this hypothesis, supporting the notion that optimal cognitive performance is characterized by brain states closer to criticality. It will be of interest to conduct critical analysis at different stages on animals with permanent cognitive impairment produced by chronic scopolamine administration which may provide valuable insights for detecting biomarkers of early-stage neurodegenerative diseases. It is worth noting that scopolamine has been found in some studies to directly affect calcium conductances, raising the question of whether the miniscope measurements might be deleteriously affected (Tsubokawa and Ross, 1997; Power and Sah, 2002). This is unlikely, as other scopolamine experiments conducted in this laboratory (Sun et al., 2021) using miniscopes, as well as electrophysiological studies (Brazhnik et al., 2004), have shown very similar effects.

In conclusion, we present for the first time to our knowledge clear evidence of shifting of neuronal ensemble behaviour closer to a critical state when engaging in a cognitive task and that conversely impairment of cognitive capability is associated with a shift away from a critical state. It will be of great interest to see if changes in critical dynamics in vivo can be used as a biomarker of cognitive dysfunction, and even perhaps a control variable for restorative interventions.

## Acknowledgments

This project was supported by the Australian Research Council’s Discovery Projects funding scheme (Project DP220101166) and the RMH Neuroscience Foundation.

## Author contributions

Conceptualization, C.F., F.H., and A.N.B.; Surgery, D.S.; Experiment, F.H., and D.S.; Methodology, F.H.; Original Draft, F.H.; Review & Editing, F.H., D.S., C.F. and A.N.B.; Supervision, C.F., and A.N.B..

## A Supplementary Materials

### A.1 Ethical Considerations

All surgical and experimental procedures were approved by the Florey Animal Ethics Committee (No. 20-060UM) and were conducted in strict accordance with the Australian Animal Welfare Committee guidelines.

### A.2 Subjects

Eight naive adult male C57BL/6 mice aged 6 weeks were housed in The Florey Institute of Neuroscience and Mental Health, Melbourne. All animals weighed 27–30 g (28.5 g ± 0.679 g) at the time of surgery and were housed individually. The facility was kept on a 12-12 hour light-dark schedule, with the lights on from 7:30 am to 7:30 pm. The mice had ad libitum access to standard chow and water.

### A.3 Stereotaxic Surgery

We aimed to use miniaturized fluorescence microscopes (miniscopes) as our means for in vivo calcium imaging from large populations of neurons in awake and behaving animals. The surgical procedures for this technique comprised two components: viral infusion and Gradient Index (GRIN) lens implantation.

#### A.3.1 Virus Infusion

A custom-made injection system was used to inject the pAAV.Syn.GCaMP6f.WPRE.SV40 virus (500 nL, viral load: 2.2×10^1^3 GC/ml; AddGene, United States) into the dorsal hippocampus (AP −2.1, ML +2.1, DV −1.7 relative to bregma) over a duration of 15 minutes. The virus was initially loaded into a capillary (with an interior diameter of 0.9 mm) created using a Sutter P-1000 electrode puller to produce a tip diameter of 20-50 µm. The open end was then sealed using silicon oil (Sigma-Aldrich, United States); The virus volume was controlled by a round brass rod (diameter: 0.8 mm; Albion Alloys, United Kingdom), connected to a 3D positioner which was fitted into the capillary. Following viral injection, the virus injector remained in place for an additional 10 minutes to allow for viral diffusion. The animal was then left to recover for one week to enable fluorophore expression.

#### A.3.2 GRIN Lens Implantation

Anchors were implanted by placing two 1 mm screws at coordinates AP +1.8, ML −2.5 and AP −2.8, ML −0.8. A small section of the skull, cantered at AP −2.1, ML +1.6, was removed using a 2 mm drill bur, and the exposed dura was carefully cleaned using fine tweezers. AA 27-gauge blunt needle was used to aspirate the cortex, exposing the vertical striations of the hippocampal fimbria. Artificial cerebrospinal fluid was used during the procedure to keep the operating field clear. The GRIN lens (0.25 pitch, #64-519, Edmund Optics) was implanted 1.35 mm deep into the tissue surface, with the most posterior point of the drilled hole (next to lambda side) serving as a reference point (Cai et al., 2016). To prevent movement of the implanted GRIN lens, cyanoacrylate glue was applied around the lens and dental cement was applied for additional support. The lens was further protected by applying fast-setting silicone adhesive (Dragon Skin Series ®, United States). Following the surgery, the animal received intraperitoneal injections of Meloxicam (2 mg/kg) daily to alleviate pain and inflammation. Additionally, enrofloxacin water (1:150 dilution, Baytril ®, United States) was provided to the animal for one week. Four weeks after the initial surgery, a metal baseplate was attached to the animal’s head to hold the miniscope in place at the optimal focal distance. It should be noted that this step is not considered a surgical procedure.

### A.4 Training

Following the surgery, the animals were handled twice a day for approximately 10 minutes during the daytime, and their weight was measured after each handling session for five days. For 3 days prior to recordings, the animals were habituated to wearing the miniscope and moving around freely with the device attached to their heads for 10 minutes per day.

### A.5 Experimental Procedures and Behavioural Tasks

All the recordings were performed during the daytime. The animal was brought into a silent recording room 30 min before the start of recording to acclimate to the surrounding environment.

The standard novel object recognition (NOR) test was performed using the protocols introduced in previous works (Leger et al., 2013). This task was conducted in a square-shaped plastic open field arena of size 33 cm × 33 cm × 20 cm. Two identical objects of the exact same color and shape were chosen for the familiarization phase. A third (novel) object consistent in height and volume but different in shape, color, and appearance was used for the testing phase. All objects were odorless, brightly colored, not chewable, and of a shape and weight that did not allow mice to climb over them or knock them over.

During habituation, the animals were allowed to explore the empty arena for 10 minutes per day and 3 days prior to the experiment day. Twenty-four hours after the last session of habituation, the NOR experiments started where animals were exposed to the familiar arena with the two identical objects placed diagonally inside the arena and at an equal distance from the walls for a 15-minute recording session (familiarization phase). After a 10-minute inter-session-interval inside the animal’s home cage, the mice were allowed to explore the open field for another 15 minutes in the presence of one familiar object and the novel object to test long-term recognition memory (testing phase). The arena was cleaned with 80% ethanol to eliminate scent clues before the first day of habituation and was kept intact during the remainder of the experiment. All the objects were also cleaned with 80% ethanol before each recording session.

On the experiment day, prior to the familiarization phase, a 15-minute recording session was performed inside the animal’s home cage as the baseline control. Custom-made miniscope acquisition software was used to record imaging frames during all the recording sessions at a sampling rate of 30 FPS. The miniscope cable was suspended over the open arena through a custom-made commutator. TopScan (TopView Analyzing System 2.0; Clever Sys Inc.), as a top-view-based behavior analysis system, was used to detect and follow animal body parts and provide accurate behavior analysis results during NOR familiarization and testing phases.

#### A.5.1 Scopolamine injection

One month after the completion of NOR experiments on the healthy animals, 2 were removed from the study due to showing signs of reduced mobility and a significant increase in the level of their struggle while mounting miniscope devices on their baseplates. The remaining 6 animals were employed in another set of repeated NOR experiments (all settings and protocols identical to those above). In the first week, the animals were injected with a dose of saline (0.9% NaCl, intraperitoneal, i.p.; 10 mL/kg body mass), 20 minutes before the familiarization phase of NOR (Tinsley et al., 2011). After 2 weeks of no experiments, the same 6 animals were injected with a dose of scopolamine (intraperitoneal, i.p.; 0.6 mg/kg body mass) 20 minutes before the familiarization phase of NOR. Scopolamine hydrobromide (Sigma-Aldrich, United States) was dissolved in sterile 0.9% saline. This was followed by the regular protocols for both phases of NOR while recording the same CA1 neuronal networks using Miniscopes.

### A.6 Data processing

The acquired data from an experimental session with any given length consists of sequential recording files each including 100 frames. The frames were all captured from the GIRN lens field of view with a frequency of 30 Hz and each frame has a size of 480 × 752 pixels.

To start with the processing pipeline, inspired by previous work and utilizing the minian package (Dong et al., 2021), we introduce Pre-Processing, Background Removal, Motion Correction, Initialization of Constrained Non-negative Matrix Factorization (CNMF), and CNMF as the prominent building blocks of our workflow.

#### A.6.1 Pre-processing

For cases where the active neuronal population did not expand the entire field of view, the region of interest was a subset in all the video frames. Next, a glow removal step was carried out by calculating a minimum projection across all frames and subtracting it from all. When the glow in the background was removed, the recordings were denoised frame by frame. Denoising was done using a median filter on each frame with a kernel size that must be chosen according to the size of the largest cells in the field of view. Usually, an effective kernel size is assumed to equal half the diameter of the largest cell present in the video file (Dong et al., 2021).

#### A.6.2 Background removal

In this step, the background in each frame (everything except the fluorescent signal of in-focus cells) was estimated and removed. For this cause, Morphological Erosion and Dilation (Raid et al., 2014) were used respectively with the window size equal to the expected size of the largest cell diameter. This was also done frame by frame. In the erosion process, each pixel was replaced by the minimum value within the filter window, and hence any bright features that appeared smaller than the window diameter were converted to zero brightness. Following this conversion, the dilation operation, to replace each pixel with the maximum value in the filter window, is basically the reverse procedure to bring back all the bright features except those completely eliminated. The morphological opening method obtains a good estimation of the background by removing the bright cells if the window size matches the expected cell diameter. This estimated background was then removed from all cells.

#### A.6.3 Motion correction

At this step, firstly, the frame with maximum projection values was selected as the base frame. Then, for all other frames, a two-dimensional cross-correlation between that frame and the base frame was calculated using a fast Fourier transform (fft). Next, the frame with the maximum correlation was chosen and its inverse value was applied as the required correction shift to all frames.

#### A.6.4 Initialization of CNMF

To utilize the Constrained Non-negative Matrix Factorization (CNMF) algorithm (Zhou et al., 2018; Kotliar et al., 2019) on our data, an initial estimation of the spatial footprints and temporal activity traces of the neural population was required. This was done using the seed selection procedure and refinement of initial seeds by signal-to-noise ratio (SNR) evaluation in the minian pipeline (Dong et al., 2021) and initializing the spatial footprint and temporal activity matrices.

#### A.6.5 CNMF

Finally, the CNMF technique to extract an accurate estimation of the spatial footprints and temporal activities of the cells, as well as the same information about the background, was performed (Zhou et al., 2018). This method employs a constrained deconvolution (CD) approach to seek the sparsest activity signal which defines the observed fluorescence intensities up to an estimated measurement noise level.

### A.7 Data analysis

We binarized the Calcium signals of each neuron in each recorded frame (i.e., 33.3 ms) based on being active or not, i.e., 1 and 0. The network activity as the sum of all recorded activity within the same time frame was then calculated. neuronal avalanches were then defined based on the network activity by applying a threshold at the 60th percentile of total network activity (Poil et al., 2012). Consistent with critical systems, our results were robust across a range of thresholds (30 to 70%).

To demonstrate the criticality of a system, we investigated the presence of the following criteria in the dynamics of our data. These criteria are all necessary conditions of a critical regimen and together can show with high confidence whether the system lies near a critical point (Beggs and Plenz, 2003; Wilting and Priesemann, 2018).

### A.8 Multiple power-laws

A critical system has units of information that show some fluctuation in their activity while maintaining a level of correlation among them. This implies that such a system is defined by scale-free dynamics and events in both spatial and temporal domains obey power-laws. Thus, events are contiguous cascades of neuronal activity rather than being limited to local bursts or huge firing of neurons consuming the network. These activities are called neural avalanches. If a system is performing near-criticality, we are looking for not just one, but multiple power-law distributions fitted to various aspects of the system dynamics in different domains (Nishimori and Ortiz, 2010).

To investigate this property in our system, we utilize the neuronal avalanches. We define the start and end points of an avalanche when the network activity crosses a threshold value (as stated above) from below and then above (Poil et al., 2012). The size of an avalanche, S, is the total number of events during the avalanche. The avalanche duration, D, is the time between threshold crossings. Similar to (Ma et al., 2019), we use maximum likelihood estimation to fit a truncated power-law to the avalanche size distribution:

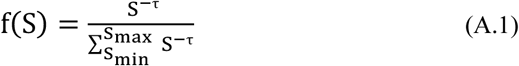

where τ would be the power-law exponent corresponding to avalanche sizes. Assuming we have detected N_A_ avalanches in the recording files, the fitting process to obtain the above equation is an iterative procedure (Klaus et al., 2011):

1. Find the maximum avalanche size S_max_.
2. Evaluate the three different power-law exponents, τ, for 3 values of minimum avalanche sizes observed, S_min_.
3. Calculate the Kolmogorov-Smirnov or KS test for this estimation to determine the goodness-of-fit between the fitted power-law and empirical distribution.
4. Among the obtained KS values, choose the smallest one alongside the corresponding τ and S_min_ values.
5. Complete the estimation if 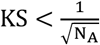 or otherwise repeat steps 2 to 5 with S_max_ reduced by 1 until this condition is met.

Steps 3 to 5 are necessary to ensure the data distribution indeed comes from a power-law rather than another candid heavy-tailed distribution such as log normal and stretched exponential forms (Clauset et al., 2009). Eventually, by applying the same procedure to the set of D values, the corresponding power-law exponent of α is calculated for avalanche duration distribution. As proposed in (Ma et al., 2019), hypothesis testing was conducted to determine if a power law fit of an avalanche distribution was plausible. To achieve this, 1000 artificial power law distributions were generated using the same power law exponent, the number of detected avalanches, and minimum and maximum avalanche sizes as the experimental distribution. These surrogate distributions were simulated using the inverse method as 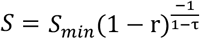 where r is a random number sampled from a uniform distribution between 0 and 1. Based on the empirical data, we upper-truncated any surrogate distribution at the maximum cutoff equivalent to S_max_. After generating the simulated surrogate distributions, the KS statistics were employed to quantify how far they are from a perfect power law. A p-value was thus determined as the ratio of the surrogate distributions with KS values smaller than those of the corresponding experimental avalanche distributions. We set the significance level to 0.05; meaning p < 0.05 implies a rejection of the power law hypothesis while p ≥ 0.05 suggests the power law hypothesis was not rejected (the fit was good).

### A.9 Exponent relation and Deviation from Criticality Coefficient (DCC)

In critical systems, there is another exponent relationship between the power law parameters (α and τ) and the exponent of mean avalanche sizes (〈*S*〉), given their duration, D (Friedman et al., 2012). We first find this third power law exponent of the system, β, from the experimental data using linear regression given the following exponent relation is present in a critical system:

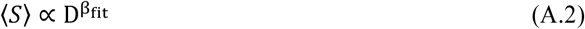

This third power law exponent also relates to the size and duration distributions of the avalanches and is predicted by:

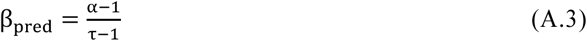

Comparing the fitted value from the empirical data (β_*fit*_) and its estimation using α and τ exponents (β_pred_), a new measure is derived to evaluate the Deviation from Criticality Coefficient (DCC), parameterized as d_CC_:

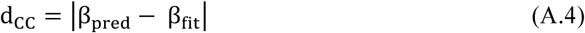

where β_pred_and β_fit_ are the predicted and fitted values of β respectively. Consequently, a smaller DCC value indicates a more accurate fit power law distribution to the empirical data.

### A.10 Scaling function

According to (Friedman et al., 2012), the obtained β can be used to accomplish a scaling function for the avalanche shapes. In a critical system, all avalanche profiles of different sizes are scaled versions of the same shape.

For any given avalanche duration D, the average number of neurons firing at time t (within D seconds) is defined by s(t, D). The following relations hold in this system:

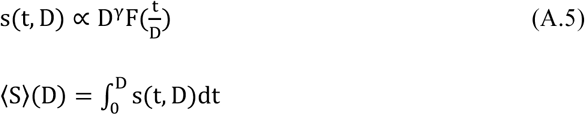

where 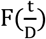 is a universal function for all avalanches and is proportional to 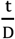 and γ = β − 1. Hence in this process, we use an initial β to predict γ, and using this γ and the first term in Equation A.5, we get 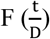 as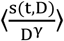. Here 〈·〉 denotes the average over all avalanches with duration D. A collection of 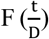functions are extracted for various D durations. The error for finding this new β is described below:

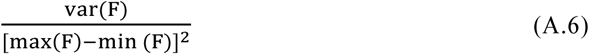

Repeating this process with various values for β, we select the exponent that produces the smallest error in Equation A.6 as the final scaling factor. So, one can measure β using both the shape collapse and the average avalanche size given a certain duration. These values must be the same when the system is truly at a critical point and hence, comparing them can help confirm the criticality idea (Marshall et al., 2016). Therefore, we expect to obtain similar (if not the same) β values from this analysis and our prediction in Section A.9, and the difference between these two β values is reported here as the SC error. We utilize the NCC toolbox in MATLAB (Marshall et al., 2016) to perform shape collapse on data. This shape collapse error is minimized under critical conditions. We used avalanches with durations between 4 and 20 bins (133.2 to 666 ms) for the shape collapse analysis. The avalanches recorded over the time course of the study were not abundant enough to conduct meaningful shape collapses beyond these cutoff points. Figure A1 illustrates the scaled avalanche shapes for the 8 animals tested in both Cage and NOR recordings.

### A.11 Branching ratio

The branching ratio is defined as the ratio of the number of neurons active at time t + 1 to the number of active neurons at time t. Since a critical regime is naturally balanced and avoids runaway gains, the branching ratio is supposed to be 1. Meaning on average, network activity neither saturates nor damps in time.

Let’s assume we have detected N active neurons in total and the number of active neurons in each time step t is defined by N(t). Considering a fixed branching ratio of m, we get 〈N(t + 1)|N(t) 〉 = mN(t) + h where 〈|〉 is the conditional expectation and h is modeling the mean rate of the external drive. The activity is stationary if m < 1, while it grows exponentially if m > 1 meaning that m = 1 separates these two regimens and represents a critical dynamic point. A precise prediction of m helps to a great extent while assessing the risk that N(t) develops a large and devastating avalanche of events such as epileptic seizures.

Under the circumstances when the full activity N(t) is known, m can be conventionally estimated using linear regression. However, under subsampling when we are aware only a fraction of neurons in a brain area is sampled, this conventional method will be biased by a certain amount. The bias vanishes only if all units are sampled because it is inherent to subsampling and cannot be overcome by obtaining longer recordings. Thus, we will instead work with the subsampled activity n(t) where the fraction of recorded units to all cells is defined as a constant µ. n(t) will be a random variable whose expectation is proportional to the real N(t) and we have < n(t)|N(t) >= µN(t) + ξ where µ and ξ are constants. According to Wilting and Priesemann (2018), the bias value for the conventional linear estimator can now be calculated as:

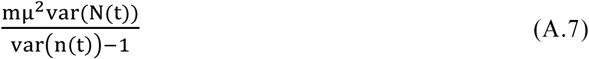

To overcome this subsampling bias, the method introduced by Wilting and Priesemann (2018) is utilized. Instead of directly using the biased regression of activity at time t and t + 1, multiple linear regressions of activity between times t and t + k are performed with different time lags k = 1, …, k_max_ = 2000. Each of these k values returns a regression coefficient r_k_ with r_1_ being equal to the result of a conventional estimator of m. Under subsampling, all these regression slopes are biased by the same factor 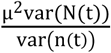. In these circumstances, instead of the exponential relation r_k_ = m^k^ which is expected under full sampling, we should generalize this equation to:

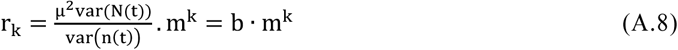

Having multiple calculated r_k_ values, we can estimate both b and m which are constant for all k.

The above multiple-regression (MR) estimator is equivalent to estimating the autocorrelation time of sub-critical periodic autoregressive models (PARs), with equal autocorrelation and regression coefficients (r_k_). While subsampling decreases the autocorrelation strength (r_k_), the decay time of the autocorrelation is indeed preserved. This intrinsic timescale allows inferring the corresponding branching parameter m of the system. To estimate this intrinsic timescale, utilizing the toolbox introduced by (Spitzner et al., 2021), we used 1) an exponential function, as well as 2) a complex function featuring an oscillatory term, a Gaussian decay term, and an offset term as decay functions fitted to r_k_ values and reported the results from the chosen function with the highest goodness of fit.

---

**Figure A1:**
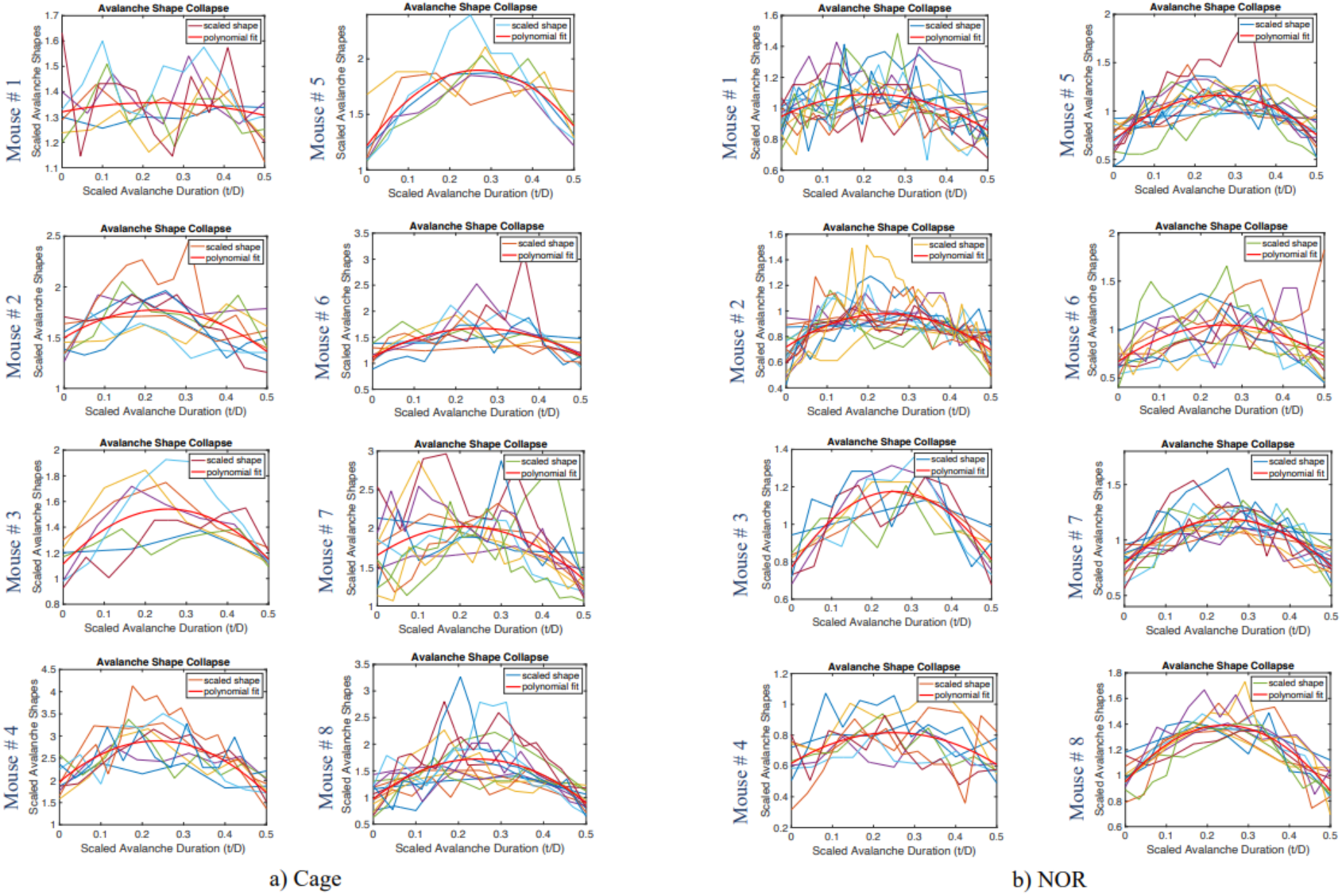
The scaled avalanche shapes across a range of durations for. **a)** Cage and **b)** NOR recordings of the 8 animals tested. These scaled shapes (in colored lines) showed smaller errors around the polynomial fit (in solid red) in the NOR recordings while this error increased significantly in the data from Cage recordings. See Figure 3.

## References

Altamura, M., Elvevåg, B., Campi, G., De Salvia, M., Marasco, D., Ricci, A. and Bellomo, A., 2012. Toward scale-free like behavior under increasing cognitive load. Complexity, 18(1), pp.38–43.

Antunes, M. and Biala, G., 2012. The novel object recognition memory: neurobiology, test procedure, and its modifications. Cognitive processing, 13, pp.93–110.

Beggs, J.M. and Plenz, D., 2003. Neuronal avalanches in neocortical circuits. Journal of neuroscience, 23(35), pp.11167–11177.

Beggs, J.M., 2022. The cortex and the critical point: understanding the power of emergence. MIT Press.

Bellay, T., Klaus, A., Seshadri, S. and Plenz, D., 2015. Irregular spiking of pyramidal neurons organizes as scale-invariant neuronal avalanches in the awake state. Elife, 4, p.e07224.

Brazhnik, E., Borgnis, R., Muller, R.U. and Fox, S.E., 2004. The effects on place cells of local scopolamine dialysis are mimicked by a mixture of two specific muscarinic antagonists. Journal of Neuroscience, 24(42), pp.9313–9323.

Cai, D.J., Aharoni, D., Shuman, T., Shobe, J., Biane, J., Song, W., Wei, B., Veshkini, M., La-Vu, M., Lou, J. and Flores, S.E., 2016. A shared neural ensemble links distinct contextual memories encoded close in time. Nature, 534(7605), pp.115–118.

Chialvo, D.R., 2010. Emergent complex neural dynamics. Nature physics, 6(10), pp.744–750.

Clancy, K.B., Koralek, A.C., Costa, R.M., Feldman, D.E. and Carmena, J.M., 2014. Volitional modulation of optically recorded calcium signals during neuroprosthetic learning. Nature neuroscience, 17(6), pp.807–809.

Clauset, A., Shalizi, C.R. and Newman, M.E., 2009. Power-law distributions in empirical data. SIAM review, 51(4), pp.661–703.

Cocchi, L., Gollo, L.L., Zalesky, A. and Breakspear, M., 2017. Criticality in the brain: A synthesis of neurobiology, models and cognition. Progress in neurobiology, 158, pp.132–152.

Deco, G. and Jirsa, V.K., 2012. Ongoing cortical activity at rest: criticality, multistability, and ghost attractors. Journal of Neuroscience, 32(10), pp.3366–3375.

Deco, G., Jirsa, V.K. and McIntosh, A.R., 2011. Emerging concepts for the dynamical organization of resting-state activity in the brain. Nature reviews neuroscience, 12(1), pp.43–56.

Dong, Z., Mau, W., Feng, Y., Pennington, Z.T., Chen, L., Zaki, Y., Rajan, K., Shuman, T., Aharoni, D. and Cai, D.J., 2022. Minian, an open-source miniscope analysis pipeline. Elife, 11, p.e70661.

Fagerholm, E.D., Lorenz, R., Scott, G., Dinov, M., Hellyer, P.J., Mirzaei, N., Leeson, C., Carmichael, D.W., Sharp, D.J., Shew, W.L. and Leech, R., 2015. Cascades and cognitive state: focused attention incurs subcritical dynamics. Journal of Neuroscience, 35(11), pp.4626–4634.

Friedman, N., Ito, S., Brinkman, B.A., Shimono, M., DeVille, R.L., Dahmen, K.A., Beggs, J.M. and Butler, T.C., 2012. Universal critical dynamics in high resolution neuronal avalanche data. Physical review letters, 108(20), p.208102.

Gautam, S.H., Hoang, T.T., McClanahan, K., Grady, S.K. and Shew, W.L., 2015. Maximizing sensory dynamic range by tuning the cortical state to criticality. PLoS computational biology, 11(12), p.e1004576.

Ghosh, K.K., Burns, L.D., Cocker, E.D., Nimmerjahn, A., Ziv, Y., Gamal, A.E. and Schnitzer, M.J., 2011. Miniaturized integration of a fluorescence microscope. Nature methods, 8(10), pp.871–878.

Gireesh, E.D. and Plenz, D., 2008. Neuronal avalanches organize as nested theta-and beta/gamma-oscillations during development of cortical layer 2/3. Proceedings of the National Academy of Sciences, 105(21), pp.7576–7581.

Habibollahi, F., Kagan, B.J., Burkitt, A.N. and French, C., 2023. Critical dynamics arise during structured information presentation within embodied in vitro neuronal networks. Nature Communications, 14(1), p.5287.

Hahn, G., Petermann, T., Havenith, M.N., Yu, S., Singer, W., Plenz, D. and Nikolić, D., 2010. Neuronal avalanches in spontaneous activity in vivo. Journal of neurophysiology, 104(6), pp.3312–3322.

Harding, E.K., Boivin, B. and Salter, M.W., 2020. Intracellular calcium responses encode action potential firing in spinal cord lamina I neurons. Journal of Neuroscience, 40(23), pp.4439–4456.

Hasselmo, M.E., 2006. The role of acetylcholine in learning and memory. Current opinion in neurobiology, 16(6), pp.710–715.

He, B.J., 2014. Scale-free brain activity: past, present, and future. Trends in cognitive sciences, 18(9), pp.480–487.

Hoffman, K.L. and McNaughton, B.L., 2002. Coordinated reactivation of distributed memory traces in primate neocortex. Science, 297(5589), pp.2070–2073.

Jannesari, M., Saeedi, A., Zare, M., Ortiz-Mantilla, S., Plenz, D. and Benasich, A.A., 2020. Stability of neuronal avalanches and long-range temporal correlations during the first year of life in human infants. Brain Structure and Function, 225, pp.1169–1183.

Klaus, A., Yu, S. and Plenz, D., 2011. Statistical analyses support power law distributions found in neuronal avalanches. PloS one, 6(5), p.e19779.

Kowalczyk, J., Kurach, Ł., Boguszewska-Czubara, A., Skalicka-Woźniak, K., Kruk-Słomka, M., Kurzepa, J., Wydrzynska-Kuźma, M., Biała, G., Skiba, A. and Budzyńska, B., 2020. Bergapten improves scopolamine-induced memory impairment in mice via cholinergic and antioxidative mechanisms. Frontiers in Neuroscience, 14, p.730.

Leger, M., Quiedeville, A., Bouet, V., Haelewyn, B., Boulouard, M., Schumann-Bard, P. and Freret, T., 2013. Object recognition test in mice. Nature protocols, 8(12), pp.2531–2537.

Ma, Z., Turrigiano, G.G., Wessel, R. and Hengen, K.B., 2019. Cortical circuit dynamics are homeostatically tuned to criticality in vivo. Neuron, 104(4), pp.655–664.

Marshall, N., Timme, N.M., Bennett, N., Ripp, M., Lautzenhiser, E. and Beggs, J.M., 2016. Analysis of power laws, shape collapses, and neural complexity: new techniques and MATLAB support via the NCC toolbox. Frontiers in physiology, 7, p.250.

Nielson, D.M., Smith, T.A., Sreekumar, V., Dennis, S. and Sederberg, P.B., 2015. Human hippocampus represents space and time during retrieval of real-world memories. Proceedings of the National Academy of Sciences, 112(35), pp.11078–11083.

Nishimori, H. and Ortiz, G., 2011. Elements of phase transitions and critical phenomena. Oxford university press.

Palutla, A., Seth, S., Ashwin, S.S. and Krishnan, M., 2023. Criticality in Alzheimer’s and healthy brains: insights from phase-ordering. Cognitive Neurodynamics, pp.1–9.

Petermann, T., Thiagarajan, T.C., Lebedev, M.A., Nicolelis, M.A., Chialvo, D.R. and Plenz, D., 2009. Spontaneous cortical activity in awake monkeys composed of neuronal avalanches. Proceedings of the National Academy of Sciences, 106(37), pp.15921–15926.

Poil, S.S., Hardstone, R., Mansvelder, H.D. and Linkenkaer-Hansen, K., 2012. Critical-state dynamics of avalanches and oscillations jointly emerge from balanced excitation/inhibition in neuronal networks. Journal of Neuroscience, 32(29), pp.9817–9823.

Ponce-Alvarez, A., Jouary, A., Privat, M., Deco, G. and Sumbre, G., 2018. Whole-brain neuronal activity displays crackling noise dynamics. Neuron, 100(6), pp.1446–1459.

Power, J.M. and Sah, P., 2002. Nuclear calcium signaling evoked by cholinergic stimulation in hippocampal CA1 pyramidal neurons. Journal of Neuroscience, 22(9), pp.3454–3462.

Priesemann, V. and Shriki, O., 2018. Can a time varying external drive give rise to apparent criticality in neural systems?. PLoS computational biology, 14(5), p.e1006081.

Priesemann, V., Valderrama, M., Wibral, M. and Le Van Quyen, M., 2013. Neuronal avalanches differ from wakefulness to deep sleep–evidence from intracranial depth recordings in humans. PLoS computational biology, 9(3), p.e1002985.

Priesemann, V., Wibral, M., Valderrama, M., Pröpper, R., Le Van Quyen, M., Geisel, T., Triesch, J., Nikolić, D. and Munk, M.H., 2014. Spike avalanches in vivo suggest a driven, slightly subcritical brain state. Frontiers in systems neuroscience, 8, p.108.

Pu, J., Gong, H., Li, X. and Luo, Q., 2013. Developing neuronal networks: self-organized criticality predicts the future. Scientific reports, 3(1), p.1081.

Raid, A.M., Khedr, W.M., El-Dosuky, M.A. and Aoud, M., 2014. Image restoration based on morphological operations. International Journal of Computer Science, Engineering and Information Technology, 4(3), pp.9–21.

Ribeiro, T.L., Copelli, M., Caixeta, F., Belchior, H., Chialvo, D.R., Nicolelis, M.A. and Ribeiro, S., 2010. Spike avalanches exhibit universal dynamics across the sleep-wake cycle. PloS one, 5(11), p.e14129.

San Tang, K., 2019. The cellular and molecular processes associated with scopolamine-induced memory deficit: A model of Alzheimer’s biomarkers. Life sciences, 233, p.116695.

Schindler, A.G., Li, S. and Chavkin, C., 2010. Behavioral stress may increase the rewarding valence of cocaine-associated cues through a dynorphin/κ-opioid receptor-mediated mechanism without affecting associative learning or memory retrieval mechanisms. Neuropsychopharmacology, 35(9), pp.1932–1942.

Sethna, J.P., Dahmen, K.A. and Myers, C.R., 2001. Crackling noise. Nature, 410(6825), pp.242–250.

Shew, W.L. and Plenz, D., 2013. The functional benefits of criticality in the cortex. The neuroscientist, 19(1), pp.88–100.

Spitzner, F.P., Dehning, J., Wilting, J., Hagemann, A., P. Neto, J., Zierenberg, J. and Priesemann, V., 2021. MR. Estimator, a toolbox to determine intrinsic timescales from subsampled spiking activity. Plos one, 16(4), p.e0249447.

Sreekumar, V., Dennis, S., Doxas, I., Zhuang, Y. and Belkin, M., 2014. The geometry and dynamics of lifelogs: discovering the organizational principles of human experience. PloS one, 9(5), p.e97166.

Stewart, C.V. and Plenz, D., 2006. Inverted-U profile of dopamine–NMDA-mediated spontaneous avalanche recurrence in superficial layers of rat prefrontal cortex. Journal of neuroscience, 26(31), pp.8148–8159.

Sun, D., Shaik, N.E.K., Unnithan, R.R. and French, C., 2024. Hippocampal cognitive and relational map paradigms explored by multisensory encoding recording with wide-field calcium imaging. Iscience, 27(1), p.108603.

Sun, D., Unnithan, R.R. and French, C., 2021. Scopolamine impairs spatial information recorded with “miniscope” calcium imaging in hippocampal place cells. Frontiers in neuroscience, 15, p.640350.

Sun, D., Yu, Y., Habibollahi, F., Unnithan, R.R. and French, C., 2023. Real-time multimodal sensory detection using widefield hippocampal calcium imaging. Communications Engineering, 2(1), p.91.

Tinsley, C.J., Fontaine-Palmer, N.S., Vincent, M., Endean, E.P., Aggleton, J.P., Brown, M.W. and Warburton, E.C., 2011. Differing time dependencies of object recognition memory impairments produced by nicotinic and muscarinic cholinergic antagonism in perirhinal cortex. Learning & Memory, 18(7), pp.484–492.

Touboul, J. and Destexhe, A., 2017. Power-law statistics and universal scaling in the absence of criticality. Physical Review E, 95(1), p.012413.

Trautmann, E.M., O’Shea, D.J., Sun, X., Marshel, J.H., Crow, A., Hsueh, B., Vesuna, S., Cofer, L., Bohner, G., Allen, W. and Kauvar, I., 2021. Dendritic calcium signals in rhesus macaque motor cortex drive an optical brain-computer interface. Nature communications, 12(1), p.3689.

Wilting, J. and Priesemann, V., 2018. Inferring collective dynamical states from widely unobserved systems. Nature communications, 9(1), p.2325.

Wilting, J. and Priesemann, V., 2019. Between perfectly critical and fully irregular: a reverberating model captures and predicts cortical spike propagation. Cerebral Cortex, 29(6), pp.2759–2770.

Wirtshafter, H.S. and Disterhoft, J.F., 2022. In vivo multi-day calcium imaging of CA1 hippocampus in freely moving rats reveals a high preponderance of place cells with consistent place fields. Journal of Neuroscience, 42(22), pp.4538–4554.

Zhou, P., Resendez, S.L., Rodriguez-Romaguera, J., Jimenez, J.C., Neufeld, S.Q., Giovannucci, A., Friedrich, J., Pnevmatikakis, E.A., Stuber, G.D., Hen, R. and Kheirbek, M.A., 2018. Efficient and accurate extraction of in vivo calcium signals from microendoscopic video data. elife, 7, p.e28728.

Zimmern, V., 2020. Why brain criticality is clinically relevant: a scoping review. Frontiers in neural circuits, 14, p.565335.

